# Voluntary modification of rapid tactile-motor responses during reaching differs from its visuo-motor counterpart

**DOI:** 10.1101/2020.04.01.020529

**Authors:** Sasha Reschechtko, J. Andrew Pruszynski

## Abstract

People commonly hold and manipulate a variety of objects in everyday life, and these objects have different physical properties. In order to successfully control this wide range of objects, people must associate new patterns of tactile stimuli with appropriate motor outputs. We performed a series of experiments investigating the extent to which people can voluntarily modify tactile-motor associations in the context of a rapid tactile-motor response guiding the hand to a moving target (previously described in Pruszynski et al. 2016) by using an anti-reach paradigm in which participants were instructed to move their hands in the opposite direction of a target jump. We compared performance to that observed when people make visually guided reaches to a moving target (cf. Day & Lyon 2000, Pisella et al. 2000). When participants had visual feedback, motor responses during the anti-reach task showed early automatic responses toward the moving target before voluntary modification to move in the instructed direction. When the same participants had only tactile feedback, however, they were able to suppress this early phase of the motor response, which occurs less than 100 ms after the target jump. Our results indicate that, while the tactile motor and visual motor systems both support rapid responses that appear similar under some conditions, these responses show sharp distinctions in terms of their malleability.

## Introduction

Many everyday actions, like reaching for a bottle’s cap while holding the bottle, require fast and precise bimanual coordination. We recently described a tactile-motor reflex — in which a tactile stimulus delivered to one hand drives short-latency online correction of reaching with the other hand — that can support such coordination (Pruszynski et al. 2016). This tactile-motor reflex appears similar to short-latency visually-guided reactions that help guide the reaching hand toward a moving target (Day and Lyon 2000, Franklin & Wolpert 2008, Goodale et al. 1986, Gu et al. 2016, Paulignan et al. 1991, Pelisson et al. 1986, Prablanc & Martin 1992, Pruszynski et al. 2010, Saijo et al. 2005).

One hallmark of short-latency visual reactions is that they are tightly coupled to the position of the visual target. When people are instructed to respond to a target jump by moving in the *opposite* direction of the jump (“anti-reach”), they exhibit a delay in production of the correct movement compared to that observed when executing a movement in the same direction of the jump (“pro-reach”). People have difficulty suppressing this sort of movement when they are instructed to do so: they first exhibit an erroneous pro-reach towards the target (Day & Lyon 2000, Gu et al. 2016, Pisella et al. 2000) similar to the erroneous pro-saccades observed at low latency during anti-saccade tasks (Fischer & Weber 1992, Gottlieb & Goldberg 1999, Everling et al. 1999). These events are attributed to the automaticity of the pro-reach and the additional time it takes to prepare a voluntary action in the opposite direction owing to the necessity of processing stimulus location and remapping this stimulus, with respect to the instruction, to a specific motor output.

While short-latency reaching corrections have been observed in response to tactile stimulus, stimulus-response mappings for tactile stimuli are not necessarily similar to those for visual stimuli. Unlike visual information, tactile information is not as directly related to the position of objects in external space. In fact, relationships between tactile information and spatial coordinates are highly variable across everyday objects, or even on the same object when it is held in different orientations. Consider using a pool cue to precisely strike the cue ball or a fly rod to cast a fishing lure to a specific location: owing to the different physical properties of the cue stick (rigid) and fly rod (flexible), maneuvering them to specific spatial locations requires mapping tactile information in vastly different ways. Here, we investigate the flexibility of this tactile-motor reflex to remap stimuli to outputs by using an anti-reach task. We show that this reflex is indeed more flexible than the classic visual stimulus-locked response in terms of muscle activation patterns; however, it is not completely malleable and it does not result in faster anti-reach kinematic responses, at least on the one-hour timescale of this study.

## Methods

### Participants

A total of 56 unique participants (18-35 years old; 34 women) participated in three experiments. One cohort of 20 individuals participated in Experiments 1a and 1b; 16 participants performed Experiment 2; and another cohort of 20 participated in Experiments 3a and 3b. All participants provided written informed consent in accordance with methods approved by the Health Sciences Research Ethics Board at Western University.

### Procedures

Participants sat at a table in front of the experimental apparatus. Each participant used their dominant hand to reach from a start position to a spherical target (4 cm diameter) mounted on a 30 cm long carbon fiber rod (Fig 1a). Participants used a finger of their nondominant hand (thumb or index finger) to feel a tactile stimulus (which varied depending on experiment) mounted in line with the rod (Fig 1b,c). A high speed stepper motor rotated the rod, stimulus and target on some trials. On trials where the target moved, its movement was triggered when the participant lifted their finger from the start position (measured with a resistive pressure sensor) to begin reaching. The latency between liftoff and the initiation of movement was measured at approximately 30 ms, and the rotation movement took 50 ms to complete. Participants received an auditory “ready” cue, after which they could initiate a reach toward the target at a self-selected time. Participants also received an auditory cue 300 ms after the target movement (or time when the target would have moved); they were instructed to try to finish reaching by the time they heard this second cue as a pacing method.

**Figure 1.**
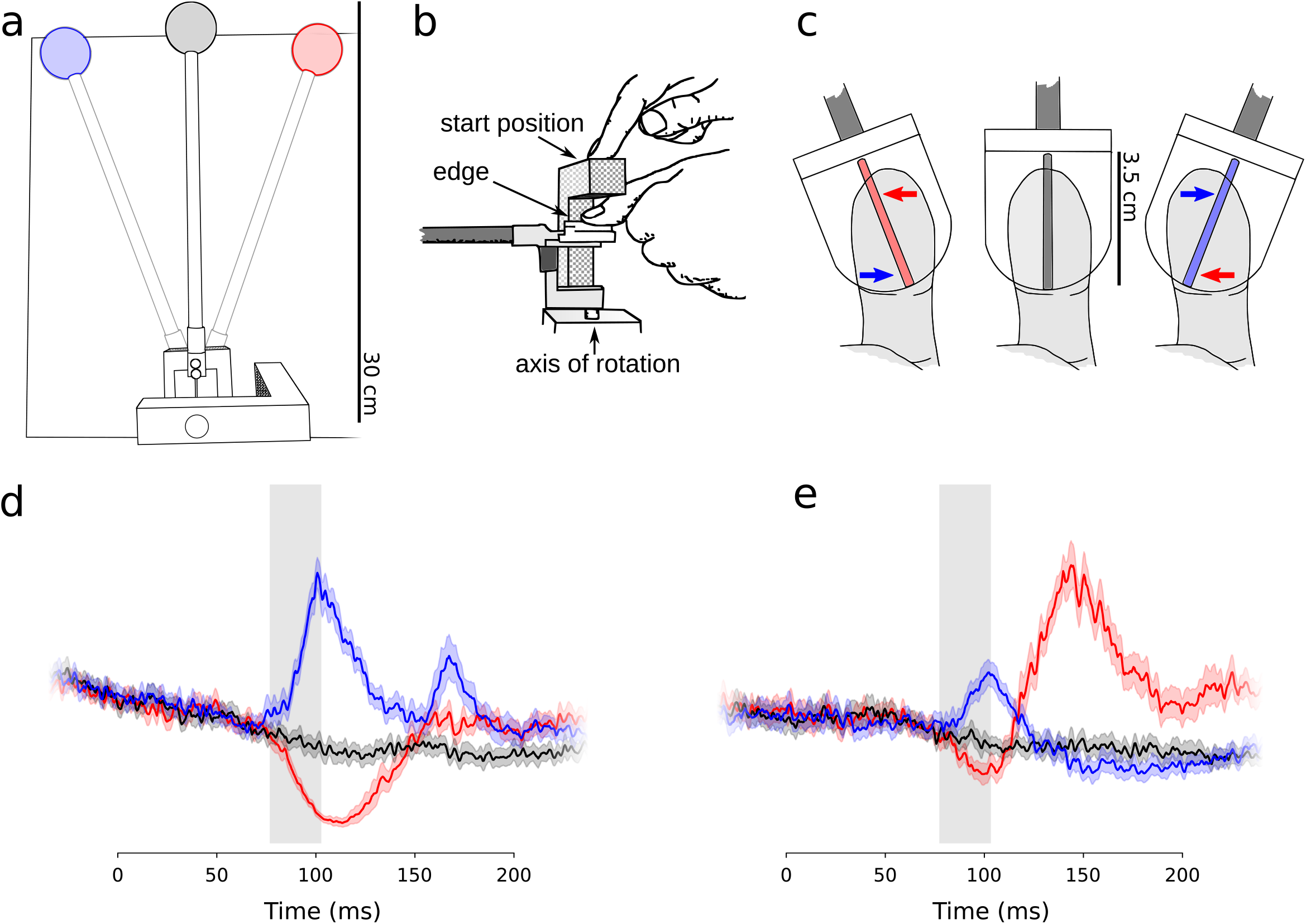
Experimental Setup. (a) Illustration of the experimental setup as viewed from the top, illustrating the position of the target in its initial position (gray), after CW jump (red), and after CCW jump (blue). (b) Illustration of participant’s starting position for all trials. (c) Illustration of the configuration of edge stimulus viewed from underneath the fingertip following CW jump (red), no jump (black), and CCW jump (blue); arrows indicate the relative motion of different parts of the edge about the center of rotation. (d, e): Across-participant mean muscle activity (EMG) traces. Thick line represents the mean and the shaded area represents the standard error of the mean. Colors represent the CW (red), CCW (blue), and no (black) jump conditions. Data aligned on rotation onset. The gray box illustrates the time during which EMG data were analyzed: a 25 ms epoch following median time of discrimination via ROC for reach instruction for a given participant (d, gray area, see Methods); the same time was then used to analyze EMG for the anti-reach instruction (e).

#### Experiment 1a: Reaching with visual and tactile feedback or tactile feedback only

Participants (n=20; 13 women; 1 left-handed) reached toward the target using either visual and tactile feedback or tactile feedback only. Tactile feedback was provided using a raised edge oriented in line with the rod holding the target ball and spanning the entire surface of the finger (Pruszynski et al. 2016). During the touch only condition, we occluded participants’ vision using LCD shutter glasses (PLATO, Translucent Technologies, Toronto, ON, CAN) which closed before participants received the ready cue and opened 300 ms after the target moved. Participants completed 240 trials, blocked according to whether visual feedback was available (120 per feedback condition); in one third of trials in each block (40), the target did not move; in the other trials, the target rotated 15° clockwise (right) or counterclockwise (left). Target movement was quasi-randomized such that all participants performed 40 trials with each movement (left, right, or none), but the target movement on a given trial was unpredictable.

#### Experiment 1b: Anti-Reaching with visual and tactile feedback or tactile feedback only

The same participants who participated in Experiment 1a also performed Experiment 1b. The procedure was identical to that of Experiment 1a, except participants were instructed to reach in the *opposite direction* of target movement using either vision and touch or touch only. Individuals participated in Experiment 1a and Experiment 1b one after the other for a total of 480 trials, but the order of participation was balanced such that half of the participants first performed Experiment 1a and the other half performed Experiment 1b first.

#### Experiment 2: Anti-Reaching with visual and tactile feedback, visual feedback only, or tactile feedback only

Another cohort of participants (n = 16; 10 women) performed an anti-reaching experiment very similar to Experiment 1b. However, whereas participants in Experiment 1b who received visual feedback also felt the edge with their nondominant finger, we added a condition in Experiment 2 in which participants did not touch anything with their nondominant finger. In each of the three feedback conditions, participants again performed 40 trials with target movements in each direction, performing a total of 360 trials during the experimental session. Participants performed all trials in a given feedback condition as a block; the order of these blocks was balanced across participants.

#### Experiment 3a: Reaching with texture tactile feedback

A third cohort of participants (n = 20; 10 women; 1 left-handed) reached to a target using visual and tactile feedback or tactile feedback only. The procedure was identical to Experiment 1 except that tactile feedback was provided with a sheet of fine sandpaper (320 grit; ~50 micron) on a flat plate, rather than a raised edge as in Experiment 1.

#### Experiment 3b: Anti-Reaching with texture tactile feedback

The participants from Experiment 3a also performed Experiment 3b. The procedure was identical to experiment 3a, except participants were instructed to reach in the *opposite direction* of target movement. Individuals participated in Experiments 3a and 3b during the same visit; half of the participants first performed Experiment 3a while the other half first performed Experiment 3b.

### Analysis

We recorded kinematics at 240 Hz using a magnetic tracker (Liberty; Polhemus, Colchester, VT, USA) attached to each participant’s dominant index finger. We recorded muscle activity using wireless electromyography (Trigno; Delsys Inc, Natick, MA, USA) with electrode modules placed on the anterior deltoid, long head of the biceps, posterior deltoid, and long head of the triceps. EMG data were digitized at 2000 Hz using a 16-bit ADC (USB-6225, National Instruments, Austin, TX, USA). Kinematics data were logged using the tracker’s native software (Pimgr), while the EMG data were logged using a program written in Python (Python Software Foundation, Beaverton, OR, USA) which also monitored the resistive pressure sensor to move the target and sent a synchronization signal to the kinematics logging software. We imposed a number of kinematic criteria on each individual trial to ensure that participants did not wait for the target to jump before moving toward the target, moved toward the initial target position rather than guessing the location to which the target would jump, and ended up reaching the target. We full-wave rectified muscle activity and filtered it using a 4th order, forward and reverse Butterworth filter with a passband of 40-250 Hz, and normalized to the average maximum value observed in that muscle over all trials in a given instruction and feedback combination.

We analyzed muscle activity in 25 ms windows defined by the median time at which muscle responses diverged to track leftward (CCW) and rightward (CW) target jumps (shaded regions in Figure 1d,e). We assessed the time of divergence for each participant individually using a receiver-operator characteristic (ROC); we define the time of divergence as the last local minimum or maximum preceding the time at which the ROC could successfully differentiate between 60% of trials. We tested a variety of ROC criteria to determine the divergence, which yielded similar temporal results. We use the assessment window computed during reach trials for anti-reach trials because we are interested in comparing the muscular response at similar times following target jump depending on the reaching instruction. While we recorded from four muscles of the upper arm, we report findings from anterior deltoid because it is a prime agonist during the reaching movement and shows direction-dependent activity when participants move to different targets in the medial-lateral plane.

From our kinematic recordings, we analyzed two aspects of kinematic behavior: correct response latency and kinematic “crossover.” To assess the time when a given trial showed a correct response (Figure 4b,c), we computed the medial-lateral velocity of the hand trajectory for every trial in which the target jumped. The time of correct response was defined as the last time the velocity trace crossed zero in the direction of the target jump (or in the opposite direction during anti-reach). To obtain crossover (Figure 6), we first computed each participant’s average position trajectory toward CW and CCW target jumps under each instruction (reach and anti-reach). We then computed crossover as the maximum width of the area enclosed by overlapping CW and CCW traces; if the traces diverged without overlapping, we define crossover as 0. We also illustrate kinematic behavior (Figure 3a,b) using the heading of the velocity vector.

We carried out paired t-tests in Python using the *ttest_rel* command in the *Scipy* library (Virtanen et al. 2020); for independent t-tests we used the *ttest_ind* command, and for correlations we used the *pearsonr* command in the same library. We ran repeated measures ANOVAs using linear mixed models analysis in R (R Core Team, 2020) with independent intercepts and individual subjects treated as random variables and reduced maximum likelihood fitting. We obtained p-values for effects using the Kenward-Roger (Kenward & Roger, 1997) approximation for denominator degrees of freedom as implemented in the packages *limerTest* (Kuznetsova et al, 2017) and *pbkrtest* (Halekoh & Højsgaard, 2014), as recommended by Luke (2017) for small sample sizes. We used the package *emmeans* (Lenth 2020) for post-hoc testing on effects more than two levels utilizing Bonferroni corrections for multiple comparisons.

## Results

Participants performed unconstrained reaches toward and away from a physical target (a 4 cm diameter ball on the end of a 30 cm rod) using their dominant hands. Participants used their non-dominant hands to feel the orientation of the rod (and therefore the location of the ball); on some trials they could see the ball (visual and tactile feedback), while on other trials they could only feel the location of the ball (tactile feedback only). Participants’ success varied depending on the instruction (reach/anti-reach) and type of feedback available. Under the reach instruction, the median number of accepted trials (out of 40) was 39 for vision and touch, and 35 for touch only. Under the anti-reach instruction, the median number of accepted trials was 34 for vision and touch and 32 for touch only. Average reach trajectories in 3D and in the anterior-posterior/medial-lateral plane for a single participant and for all participants together are shown in Figure 2.

**Figure 2.**
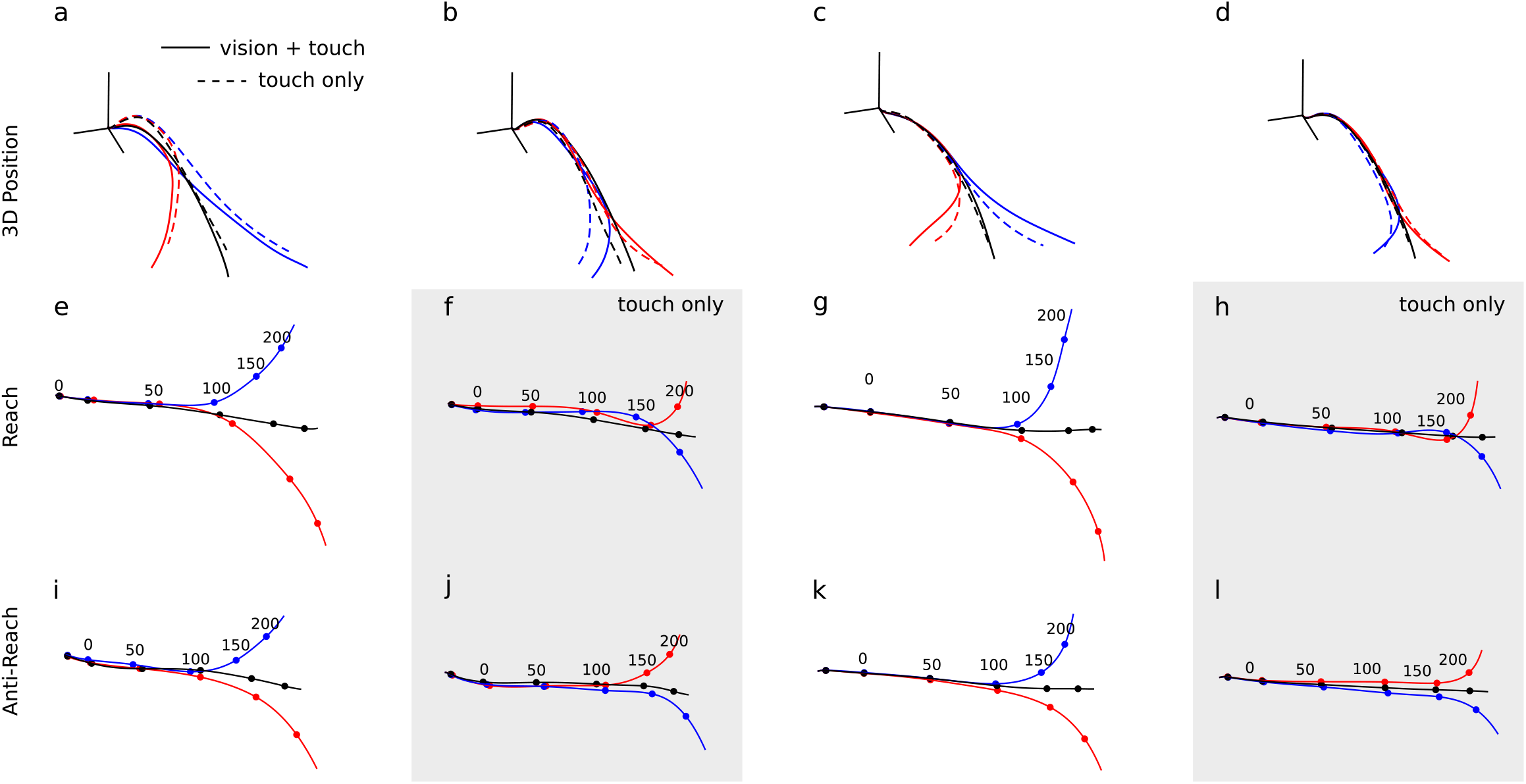
Basic Behavioral Analysis. Average kinematic behavior of a single participant (left two columns) and across-participant averages (right two columns) during Experiment 1. Markers provide timing information (in ms) relative to target jump. (a, c) 3D average traces for CCW (blue), no jump (black), and CW (red) target jumps under the pro-reach instruction, solid lines from vision and touch trials, while dotted lines are from touch-only trials. (b,d) 3D average traces from the under the anti-reach experiment (e-1) average position in the anterior-posterior/ medial-lateral plane for vision and touch trials under reach (e, g) and anti-reach (f, h) and touch only trials under reach (i, k) and anti-reach instruction (j, I).

Participants moved somewhat faster when they had visual and tactile feedback than when they had tactile feedback only. Under the reach instruction, participants accelerated to an average 3D instantaneous velocity 49.6 ± 7.7 cm/s (mean ± SEM) by the time the target jump occurred and attained an average movement velocity of 112.0 ± 2.4 cm/s over the 300 ms following the target jump; when they did not have visual feedback, they were moving at an average of 37.4 ± 5.2 cm/s when the target jump occurred and moved at an average of 104.2 ± 3.0 cm/s over the next 300ms. Under the anti-reach instruction, participants with visual feedback were moving at an average of 43.3 ± 6.8 cm/s when the target jumped and moved at 100.0 ± 4.1 cm/s over the next 300 ms, whereas participants with only tactile feedback moved averaged 39.7 ± 6.0 cm/s at target jump and 103.6 ± 3.4 cm/s over the next 300 ms. The aforementioned velocities are all in 3D; in the anterior-posterior direction only, the difference in speed between feedback conditions was marginal under the reach instruction when the target jumped (T_19_ = 2.19; P = 0.04), and there was no consistent difference over the 300ms following target jump (T_19_ = 1.33; P = 0.2). Under the anti-reach instruction, there was no consistent difference in average instantaneous speed between feedback conditions during either analysis epoch.

### Reaches toward a jumping target are rapidly updated via tactile information

Under the reach instruction, when the target moved during an ongoing reach, participants rapidly updated their reaches when the target jumped (Figure 3a). We used a receiver-operator characteristic (ROC) technique to find a time for each participant when muscle activity in anterior deltoid for rightward (CW) and leftward (CCW) target jumps diverged from one another. The median time of divergence when participants had visual and tactile information was 77 ms and it was 93 ms (Figure 4a, solid lines) when they only had tactile feedback; the divergence time was consistently faster in the former versus the latter as assessed by paired t-test (*T*_19_ = 4.14; P = 5.5e^−4^). The median time of correct response across all reach trials with a target jump under vision and touch was 138 ms while the time of correct response under touch only was 175 ms (Figure 4b, solid lines). EMG activity over the 25 ms following median time of divergence was modulated according to direction for both vision and touch trials (one-way ANOVA *Direction*: F_2,38_ = 83.76; P = 1.18e^−14^) and trials with only tactile feedback (F_2,38_ = 27.56; P = 4.02e^−8^); under touch, EMG activity was different for each possible target movement outcome (CW, CCW, no movement), whereas there for touch only there was no significant difference between CW and no movement responses during this analysis epoch. We subsequently compared vision+touch and touch only EMG activity by assessing the average difference in EMG activity between CW and CCW reaches during 25 ms bins beginning with the median onset time. The difference between CW and CCW EMG activity was greater for vision+touch compared to vision (paired t-test; *T*_19_= 3.97; P = 8.24e^−4^). Participants who showed more EMG activity under visual and tactile feedback across all target directions tended to show more EMG with only tactile feedback as well (r = 0.76; P = 1.4e^−12^; Figure 5a). Overall, the onset of corrections with only tactile feedback was marginally slower than reported by Pruszynski and colleagues (2016) and consistent with other reports of automatic visually-guided behaviors (Day & Lyon 2000, Gu et al 2016, Franklin & Wolpert 2008, Veerman et al 2008).

**Figure 3.**
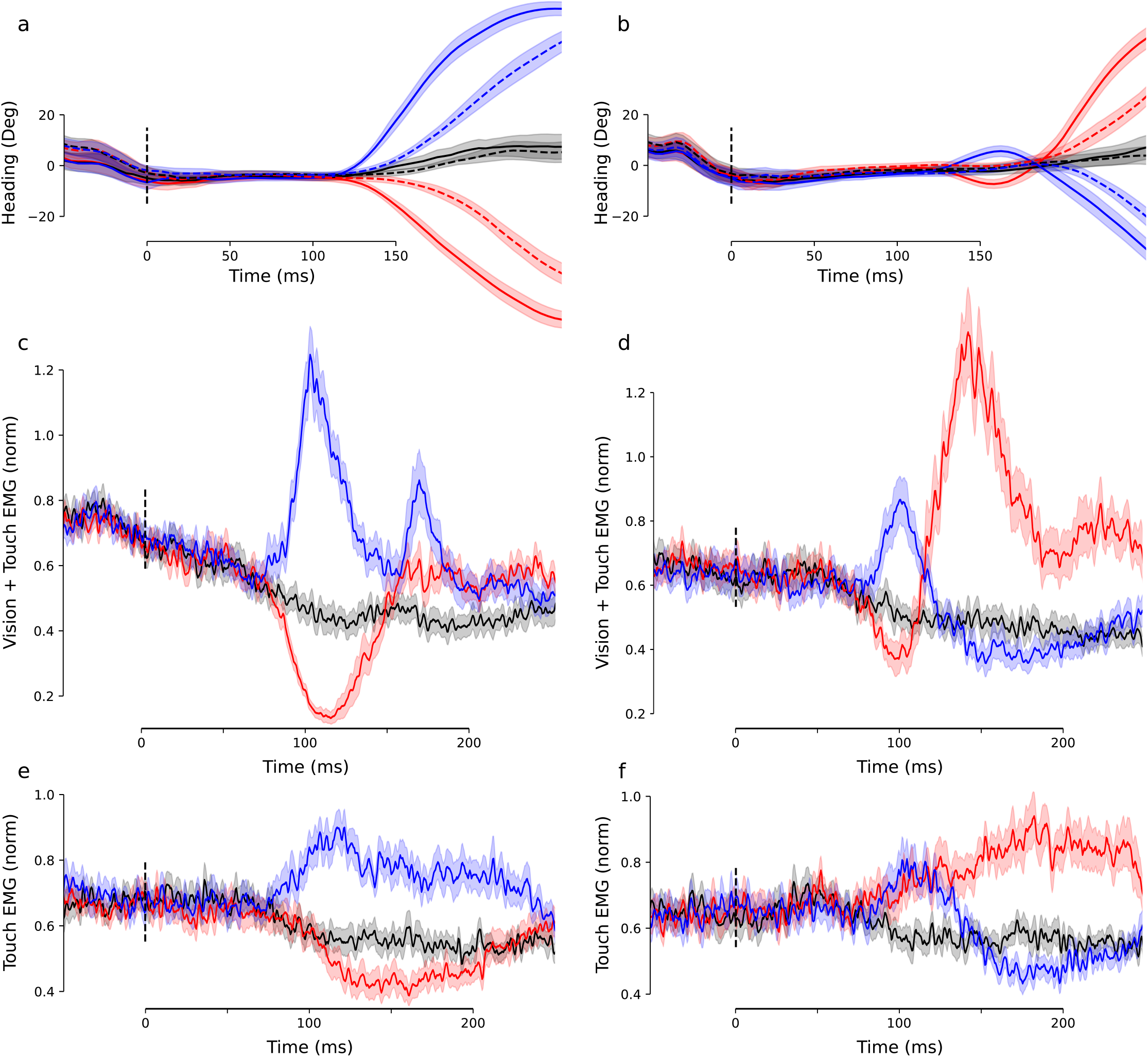
Muscle Activity in Experiment 1. (a) Average across-participant heading angle during Experiment la (pro-reach) for vision and touch (solid lines) and touch only (dotted lines). Color represents the target jump condition as in Figure 1. Shaded areas represent the standard error of the mean. (b) Average across-participants heading angle during Experiment lb (anti-reach) for vision and touch (solid lines) and touch only (dotted lines). (c) Muscle activation during vision and touch trials for pro-reach. (d) Muscle activation during vision and touch trials for vision and touch trials for anti-reach. (e) Muscle activation during touch only trials for pro-reach; (f) muscle activation during touch only trials for anti-reach.

**Figure 4.**
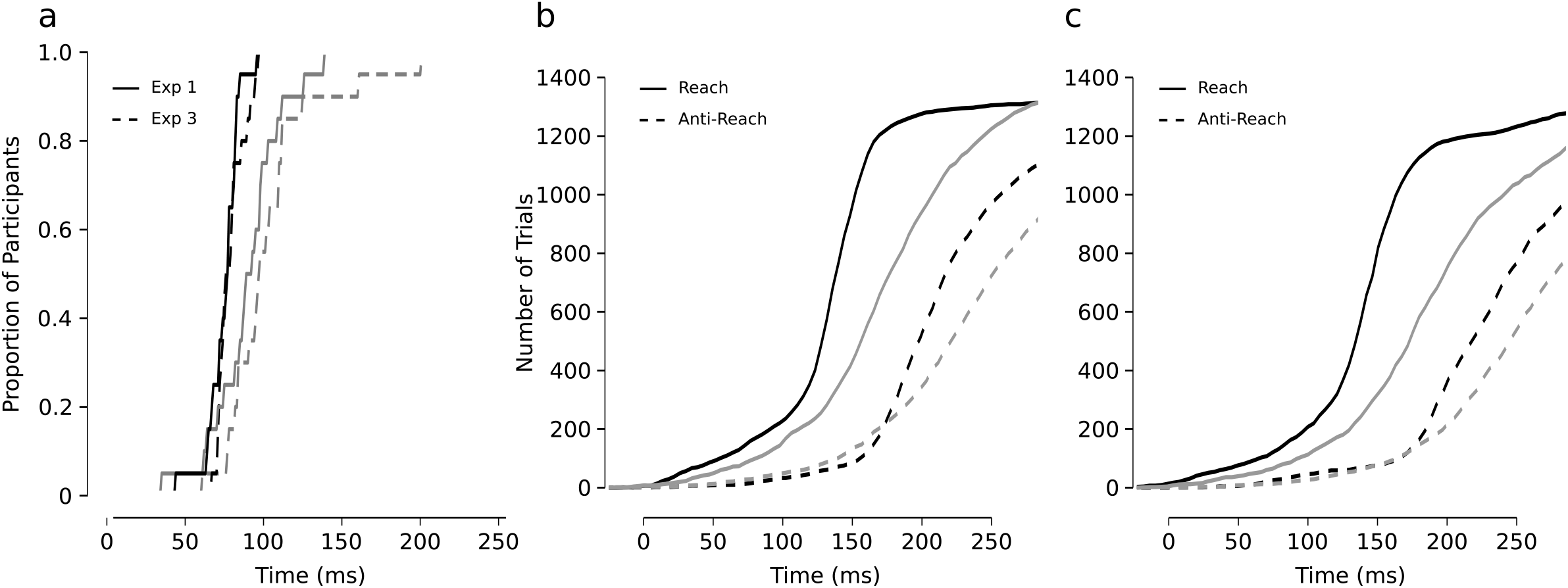
Analysis of Divergence Times. (a) Times of divergence (see Methods) for anterior deltoid EMG activity in CW and CCW trials. Solid lines represent Experiment 1 (edge stimulus); dashed lines represent Experiment 3 (texture stimulus). Black lines are for vision and touch trials, gray lines are for touch only trials. (b and c) Cumulative number of trials in Experiment 1 (b) and Experiment 3 (c) showing correct kinematic response at a given time following target movement; vision+touch trials are black, touch only trials are gray, dashed lines represent trials under the anti-reach instruction and solid lines represent trials under the reach instruction. The total number of trials shown in b and c is truncated compared to the number of trials analyzed because the onset of correct kinematic response could not be algorithmically identified in all trials.

**Figure 5.**
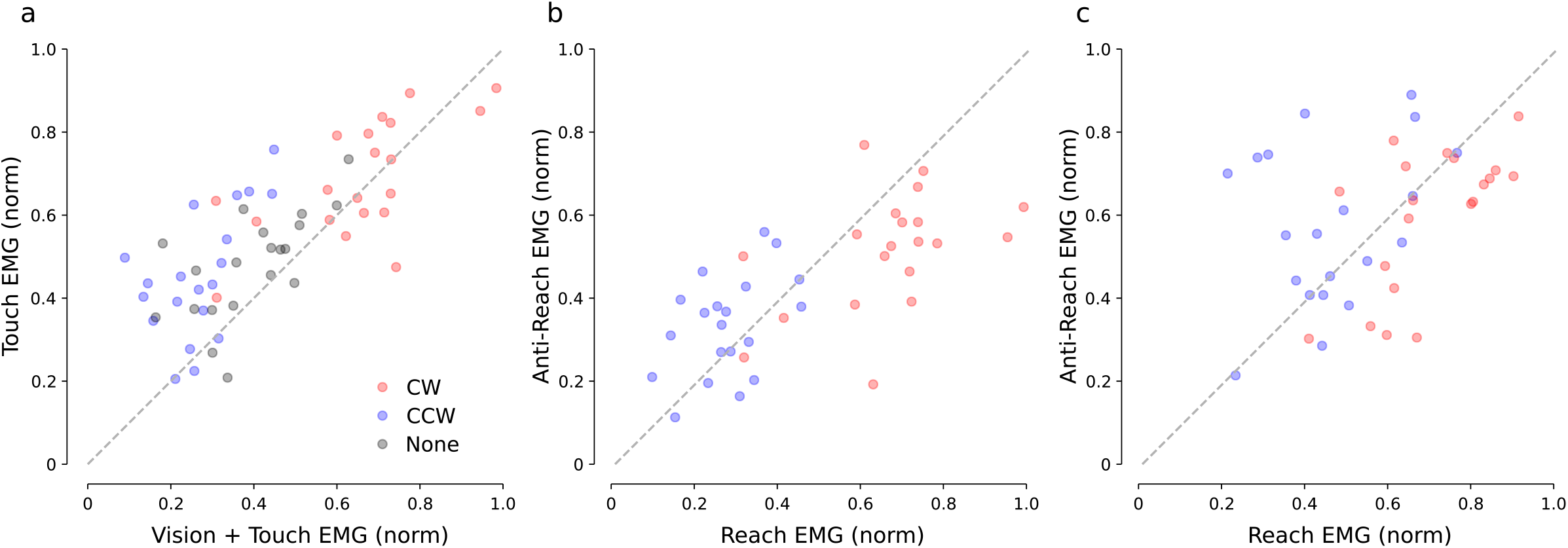
Comparison of Muscle Activity Across Feedback and Instruction in Experiment 1. (a) Average magnitude of EMG activity under the reach instruction for each participant in trials with visual and tactile feedback (x-axis) and tactile feedback only (y-axis). (b) Average magnitude of EMG activity for each participant in trials under the reach instruction (x-axis) and under the anti-reach instruction (y-axis) for trials with visual and tactile feedback. (c) Average magnitude of EMG activity for each participant under the reach instruction (x-axis) and anti-reach instruction (y-axis) for trials with tactile feedback only. For all panels, markers are color coded according to direction of target jump. Trials with no target jump (gray markers) are included only in (a).

### Movement toward an anti-reach target is suppressed but early EMG activity is evident

When the same 20 participants performed anti-reach movements with visual and tactile feedback, they made initial movements in the direction of the target jump (consistent with classical stimulus-locked response) before moving in the instructed direction. In contrast, when participants had only tactile feedback, they generally did not make these initial incorrect movements before reaching in the instructed (anti) direction (Figure 2f,h; Figure 3b). We analyzed the extent to which anti reach trajectories to CW target jumps overlapped with trajectories to CCW target jumps showed that this overlap was significantly larger during anti reach than reach for visual and tactile feedback trials (T_19_ = 5.04; P = 7.3e^−5^); this finding indicates an initial reach in the incorrect direction (see Figure 2f,h for examples of this behavior). In contrast, the instruction did not significantly modulate crossover for trials with only tactile feedback (T_19_ = 0.55; P = 0.59). The crossover observed during anti-reach was larger for visual and tactile trials compared to tactile feedback only trials (T_19_ = 4.57; P = 2.1e^−4^), as seen in Figure 6a.

**Figure 6.**
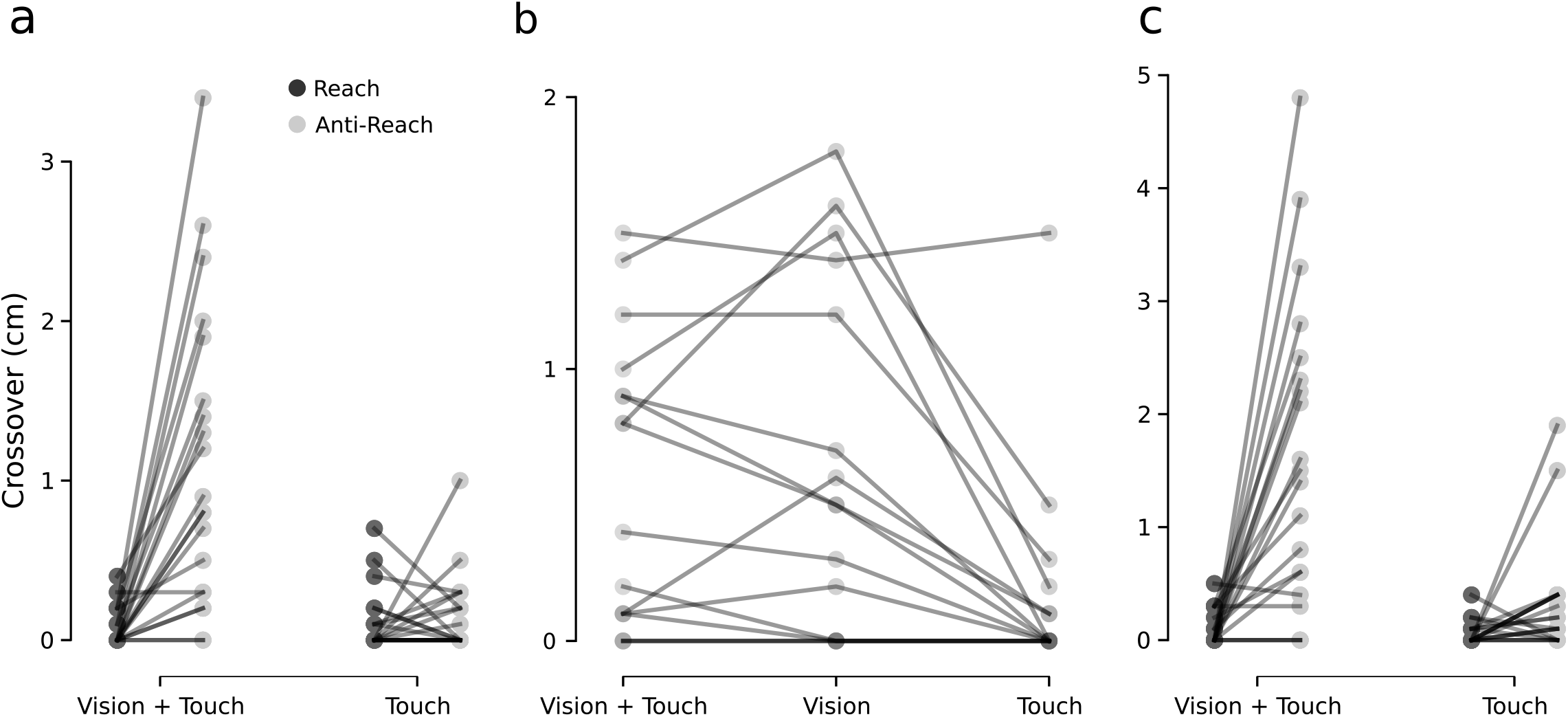
Kinematic Crossover. Maximum overlap between average trajectories to CW and CCW target jumps (in cm) for each participant, compared between feedback modalities, in Experiment 1 (a), Experiment 2 (b), and Experiment 3 (c). Dark markers represent the reach instruction; light markers represent the anti-reach instruction. Participants performed only anti-reach trials in Experiment 2.

Importantly, although participants did not move in the incorrect direction when they had only tactile feedback, their movements in the correct direction were delayed with respect to the timing observed under the pro-reach instruction. Across all trials, median time of the correct response increased from 138 ms to 208 ms with visual and tactile feedback, and from 175 ms to 229 ms for tactile feedback only (Figure 4b).

To understand the patterns of muscle activation underlying this behavior we analyzed anterior deltoid EMG in anti-reach trials in a 25 ms bin following the median divergence time in reach trials (Figure 1e). When participants had only tactile feedback, a one-way ANOVA showed a significant effect of *Direction* of target jump (F_2,38_ = 15.18; P = 1.43e^−5^). Post-hoc tests revealed elevated EMG activity compared to no jump trials for both CW and CCW target jumps, whereas there was no significant difference between EMG activity for CW and CCW target jumps at this latency, indicating a change in the early tactile response from direction-specific in reach trials to non-direction-specific in anti-reach (Figure 3f, 7d). The early response in trials with visual and tactile feedback remained direction specific, however, as shown by post-hoc tests on one-way ANOVA *Direction* (F_2,38_ = 15.18; P = 1.43e^−5^; Figure 3e, 7b), even though this results in an initial movement in the incorrect direction during the early response phase. We tested the difference between CW and CCW muscle activation using t-tests against zero for each feedback condition; although the difference was greater than zero for vision and touch (T_19_ = 6.39; P = 3.94e^−6^), indicating direction-specificity, the difference was not different from zero for touch only (T_19_ = 0.67; P = 0.51).

**Figure 7.**
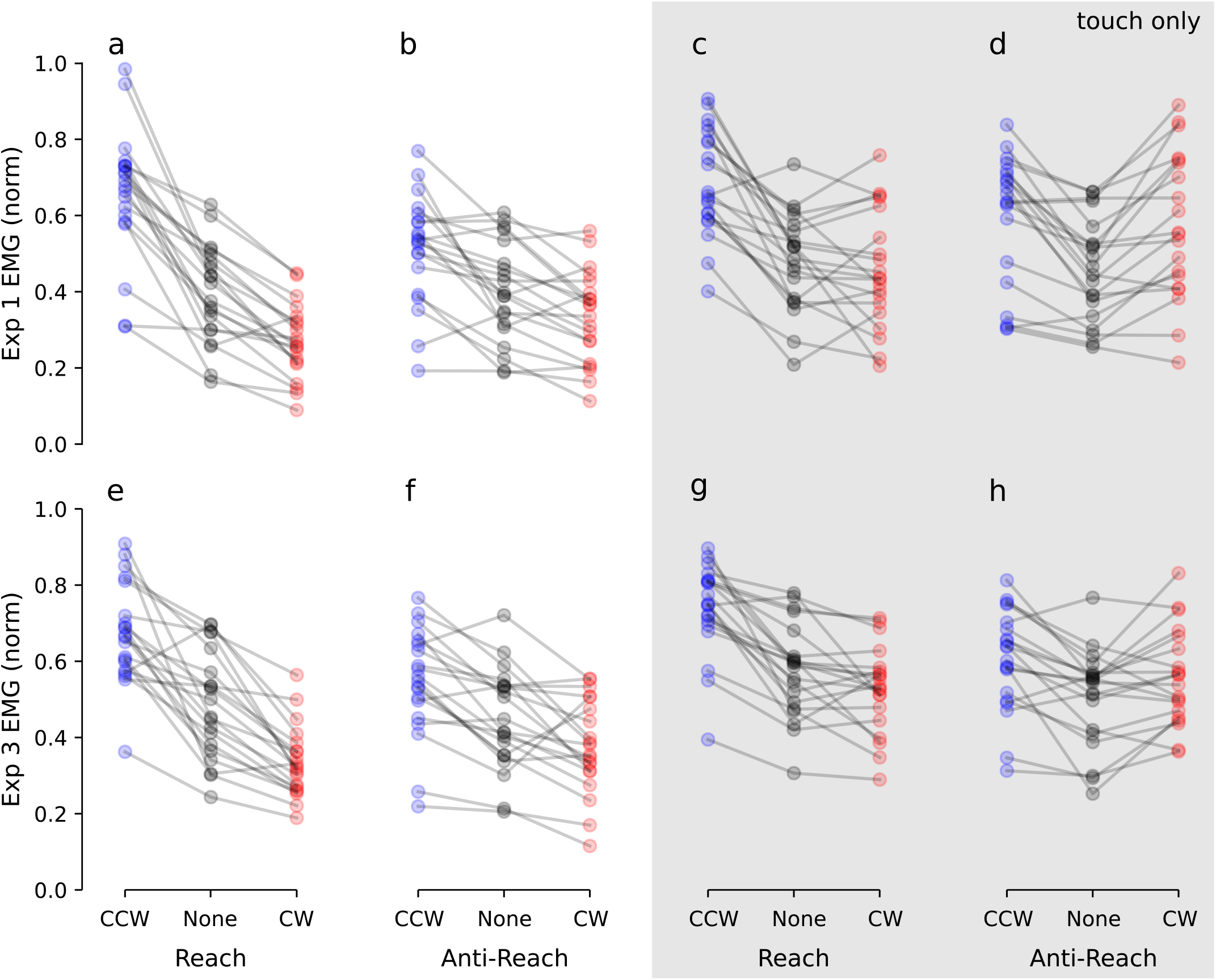
Average EMG Activity in 25 ms Window Following Divergence Time. (a-d) Average EMG activity in Experiment 1 (using an edge stimulus); (e-h) Average EMG activity in Experiment 3 (using a texture stimulus). (a,e) Average EMG activity under the reach instruction with visual and tactile feedback. (b,f) Average EMG activity under the anti-reach instruction with visual and tactile feedback. (c,g) Average EMG activity under the reach instruction with only tactile feedback. (d,h) Average EMG activity under the anti-reach instruction with only tactile feedback. Marker colors refer to direction of target jump.

Finally, we checked whether participants who showed larger EMG responses during pro-reach movements showed larger EMG responses during anti-reach movements. Under visual and tactile feedback (Figure 5b), reaches to CW target jumps were positively correlated with anti-reaches to CW target jumps (r = 0.46; P = 0.042) and reaches to CCW target jumps were positively correlated with anti-reaches to CCW target jumps (r = 0.46; P = 0.045), indicating the initial incorrect component of anti-reaches were related to behavior during the reaches when participants had visual and tactile feedback. When participants had tactile feedback only (Figure 5c), EMG during reach movements to CCW target jumps were positively correlated to anti-reach movements to CCW target jumps (r = 0.63; P = 0.003) but EMG during reach movements to CW target jumps showed no significant correlation to anti-reach movements toward CW target jumps (r = 0.31; P = 0.19) as a result of the non-direction-specific EMG response during anti-reaches with tactile feedback only.

### Vision and touch do not interfere during anti-reach

Given the differing responses to anti-reach between visual and tactile feedback and tactile feedback only, we tested whether vision and touch were interfering with one another during anti-reach. If this were the case, we would expect the early movement in the incorrect direction to be even larger when participants had only visual feedback. We recruited an additional 16 participants to perform anti-reach tasks using vision only, vision and touch (as in Experiment 1), and only touch. We observed qualitatively similar kinematic responses to the previous participants in both touch only and vision and touch conditions. There were no obvious differences in crossover between trials with visual feedback only and those with visual and tactile feedback (Figure 6b). We analyzed the difference in anterior deltoid EMG between CW and CCW target jumps for all three feedback conditions using the same 25 ms analysis windows used during pro-reaches in Experiment 1 for visual and tactile (for vision only and vision + touch data) or tactile only (for touch only) using a one-way ANOVA on *Feedback*. This test was significant (F_2,30_ = 15.31; P = 2.62e^−5^) but post hoc analyses showed that both visual and visual and tactile feedback trials were different from tactile feedback trials, while visual feedback trials were not significantly different from visual and tactile feedback trials.

### The tactile-motor reflex does not depend on persistent orientation information

The previous experiments above and in Pruszynski et aln (2016) used an edge spanning the finger pad as a tactile stimulus. The edge provides information about the position of the target both while it is moving and during steady-state because the edge remains in line with the target at the end of the movement (Figure 1c). We therefore tested whether this persistent steady-state information was necessary to engage this cross-effector tactile-motor reflex by replacing the edge stimulus with a sandpaper sheet (320 grit; average particle size < 50 μm). When participants use the sandpaper, they receive a cue over the 50 ms when the target is moving but no information about the location of the target after the target stops moving.

We recruited 20 participants (who had not participated in Experiments 1 or 2) to perform reaching and anti-reaching tasks using visual and tactile feedback or tactile feedback only. Experiments were methodologically identical to the first experiment except the tactile feedback was modified as previously described. When participants performed the pro-reach, we found median divergence times for EMG activity which were similar to those observed in Experiment 1: 76.8 ms when participants had visual and tactile feedback, and 96.5 ms when they had tactile feedback only (Figure 4a; dashed lines). During the reach instruction, participants showed EMG responses that were specific to the direction of the target jump (one-way ANOVA *Direction* for visual and tactile: F_2,38_ = 60.15; P = 1.68e^−12^; for tactile only: F_2,38_ = 28.79; P = 2.44e^−8^; Figure 7e,g). As in experiment 1, post-hoc tests indicated that there was no consistent difference between CW and NM trials for tactile feedback only, while all directions were significantly different from each other with visual and tactile feedback. Comparing the differences in muscle activity for CW and CCW target jumps revealed a larger difference in vision and touch trials than in touch only trials (T_19_ = 3.54; P = 0.0022).

When participants were instructed to perform anti-reaches with the new tactile stimulus, they again showed the kinematic patterns we saw previously. Using both visual and tactile cues, participants made early movements toward the target before moving in the instructed direction away from the target. When they had only tactile stimuli, however, early movements toward the target were not evident although they again showed a delayed reaching response compared to the pro-reach (Figure 4c). When participants had both visual and tactile feedback, the median time it took to produce a movement in the correct direction across all trials increased from 146 ms to 225 ms, and when they had tactile feedback only, median time increased from 192 ms to 250 ms. When participants only had the tactile stimulus, we still observed EMG activity in the early epoch defined by the pro-reach divergence (one-way ANOVA: F_2,38_ = 6.038; P = 0.0053). Post hoc tests indicated that EMG response did not differ during this epoch between CW and CCW target jumps; while CCW target jump EMG was different from no jump, the difference between CW and no jump was also not significant (Figure 7h). The difference in EMG activity between CW and CCW target jumps was significantly different from zero for tactile and visual feedback (T_19_ = 5.49; P = 2.68e^−5^) but it was not significantly different from zero for tactile feedback only (T_19_ = 1.52; P = 0.15).

## Discussion

The study of tactile sensation is heavily influenced by our knowledge of the vision. The tactile and visual systems are capable of extracting similar stimulus features (Pack & Bensmaia 2015), and parallels between visual and tactile processing yield interesting results (Pei et al. 2008, Pruszynski et al. 2018, Hsu et al. 2019). Our results caution against general inferences about the properties of the tactile motor system based on the visual motor system. That is, we show that although tactile and visual stimuli elicit rapid motor responses which appear similar at the outset (Prusyznski et al. 2016) they are categorically different in their susceptibility to voluntary modification.

We had participants perform reaching movements to a physical target, making online corrections when the target moved during their reach using either tactile feedback provided by an edge under their non-reaching thumb or index finger, or visual feedback in addition to tactile feedback. We also instructed the same participants to reach in the opposite direction of the target movement to perform a classical “anti-reach” paradigm. When participants had access to visual feedback during the anti-reach, they erroneously initiated movements in the direction of the target jump — consistent with previous studies (Day & Lyon 2000, Gu et al. 2016, Pisella et al. 2000) which suggest that there is something automatic and difficult to suppress about such action (Goodale et al. 1986, Paulignan et al. 1991, Pelisson et al. 1986, Prablanc & Martin 1992, Saijo et al. 2005, Veerman et al. 2008). When participants made reaches with tactile feedback only, however, they usually did not initiate movement in the wrong direction. Although tactile feedback did not result in earlier onset of movement in the instructed direction, these kinematic changes occurred as a result of qualitative changes in muscle activation patterns at low latencies (< 100 ms) following target movement. Both of these results held when another cohort of participants used a rotating texture stimulus rather than the oriented edge.

Based on the heterogeneity of tactile-motor interactions present in everyday behavior, we predicted that this reflex could be remapped during an anti-reach instruction to react as quickly as observed during a normal reach towards the new target location (i.e. pro-reach). In contrast to our predictions, rapid reaching corrections elicited by a tactile stimulus are not entirely flexible. During anti-reach tasks using tactile stimuli, kinematic corrections play out on a timescale similar to that observed in visually-guided anti-reaching. In contrast to these visually guided reaches which show minimal malleability in early responses during anti-reach, however, when participants performed anti-reaches with tactile feedback their early muscle activity changed from a pattern of agonist excitation and antagonist inhibition (as in reaching toward the target) to generalized excitation.

### The tactile-motor reflex relies on relative motion cues rather than edge orientation

The tactile-motor reflex previously we described (Pruszynski et al. 2016) could be supported by multiple features extracted early in the periphery. One possibility is that this reflex is based directly on recognizing the orientation of edges, which can be extracted very early in the tactile periphery (Pruszynski & Johansson 2014). The edge used in Experiments 1 and 2 is long enough that it could be harnessed for rapid voluntary movements with high angular precision to drive the behavioral response (as seen in Pruszynski et al. 2018). If this were the mechanism used, the anti-reach response could also be remapped at the periphery (rather than requiring a coordinate transformation for the response) by attending to only one part of the edge: in Experiments 1 and 2, the center of rotation of the edge moving under the fingertip was in the middle of the fingertip, so some parts of the stimulus (those on the finger pad distal to the center of rotation) moved in the same direction as the target, while others (proximal to the center of rotation) moved in the opposite direction (Figure 1c). If the CNS were able to selectively attend to these patterns of tactile stimulation selectively, it would provide a mechanism by which the tactile-motor reflex could rapidly implement changes in sensorimotor transformations via unambiguous tactile input.

Another possibility is that the tactile-motor reflex relies on relative motion cues, similar to the low-latency responses observed when objects slip under the fingertips (Johansson & Westling 1988). Such responses can be directionally tuned (Hager-Ross et al. 1996) and are observed bimanually and across fingers in some tasks (Cole et al. 1984, Ohki & Johansson 1999). If this were the case, remapping sensory inputs to behavioral outputs, as required in the anti-reach task, is more likely to require higher executive function similar to that posited as the reason for delayed responses in anti-saccade tasks. Our results suggest that this latter mechanism — utilizing relative motion cues and subsequently requiring coordinate transformation — is more likely to underlie the tactile-motor reflex. The most convincing reason to favor this explanation is Experiment 3, in which we show that an edge is unnecessary to elicit the tactile-motor reflex and that similar reach and anti-reach behavior are elicited when participants receive only a relative motion cue. While this leaves open the possibility that the tactile motor reflex can use edges when they are available, little appears to change in terms of response latency when participants receive only the motion cue (Figure 4). Further, the edge extraction hypothesis would suggest that remapping of the anti-reach behavior should be faster than we observed (because the relevant part of the edge stimulus could be extracted immediately), potentially being as fast as pro-reach. In contrast to this hypothesis, we observed delayed anti-reach behavior when participants had tactile feedback only. Our experiments do not speak to whether the tactile-motor reflex *could* be elicited via edge orientation alone; this would require the delivery of edge orientation without relative motion, potentially using a tactile matrix display.

### Flexibility of touch-guided sensorimotor associations

When participants performed an anti-reach task using touch, they were not able to immediately reverse their pro-reaching behavior to reach in the opposite direction at a similar latency. However, unlike anti-reaching with visual feedback, participants showed categorically different muscle responses under the anti-reach instruction. Instead of the direction-specific activation of agonist musculature and inhibition of antagonists they showed during the reach, participants showed a global increase of muscle activity during anti-reach with tactile feedback only. This contrasts with the muted direction-specific response toward the target which participants showed during anti-reach with visual and visual and tactile feedback (similar to that observed by Day & Lyon 2000). The behavior we observed when participants had access only to touch could be a building block of flexible motor responses to tactile stimuli, and indicates that the tactile motor reflex is more malleable than visually guided updates to reaching.

The limited short-term flexibility we observe in associating tactile stimuli with motor outputs could be related to understanding the dynamics of the system linking tactile stimuli to those outputs. Baugh and colleagues (2012) showed that reaction-time responses to a visuo-motor rotation are lowered — and the initial direction of the correction is more frequently correct — if a virtual tool (a pivoting link) is shown to link a participant’s hand position to a cursor. The mapping between tactile stimulus and motor output is highly variable as we use different objects or even readjust our grip on the same object. As such, a relatively safe “default” response to novel sensorimotor association involving tactile stimulus could be increased generalized muscle activity. This is consistent with responses to rapid changes in fingertip loading (Johansson & Westling 1988), which is observed across hands if both are active in the task (Ohki & Johansson 1999) and across some uninvolved fingers (Cole et al. 1984). This generalized muscle response could then be further reshaped as we learn the dynamics of the object we are handling. Further study of whether (and how completely) new tactile-motor associations can be developed over prolonged training and whether they induce after-effects could help elucidate how tactile-motor associations are built and retained.

### Limitations

One difficulty in comparing tactile-motor and visuo-motor behaviors is that it is unclear how to match the salience of tactile and visual stimuli. Our experiments made no concerted effort to match salience and participants generally reported more confidence in their abilities when they had visual feedback and we observed no difference in Experiment 2 when participants had only visual feedback compared to when they had visual and tactile feedback. In addition, while our results in Experiment 3 indicate that relative motion is sufficient to evoke this tactile-motor reflex, they do not speak to whether it is necessary as we did not apply an oriented edge stimulus without relative motion. Participants also performed all experiments with complete knowledge of how the movement of the tactile stimulus was related to target movement. It is possible that the tactile motor reflex could be more completely remapped if participants did not know how the stimulus and target movement were related (e.g. by hiding the apparatus). Similarly, participants might be able to adapt more quickly or completely to a real machine that dissociates stimulus and target position (for example using a hinge). Finally, our experiment did not test the effect of prolonged training on the remapping of the tactile-motor reflex.

## Acknowledgements

We thank Phoung Nguyen and Cynthiya Gnanaseelan for assistance with participant recruitment and data collection. In addition to the Python and R packages previously cited, we thank the developers of Inkscape (inkscape.org), as well as those of NumPy (Oliphant, 2006), Matplotlib (Hunter, 2007), and Pandas (Mckinney, 2010).

## References

Baugh LA, Hoe E, Flanagan JR. Hand-held tools with complex kinematics are efficiently incorporated into movement planning and online control. J Neurophysiol 108: 1954–1964. 2012.

Cole KJ, Gracco VL, Abbs JH. Autogenic and nonautogenic sensorimotor actions in the control of multiarticulate hand movements. Exp Brain Res 56: 582–585. 1984.

Day BL, Lyon IN. Voluntary modification of automatic arm movements evoked by motion of a visual target. Exp Brain Res 130: 159–168. 2000.

Everling S, Dorris MC, Klein RM, Munoz DP. Role of primate superior colliculus in preparation and execution of anti-saccades and pro-saccades. J Neurosci 19: 2740–2754. 1999.

Fischer B, Weber H. Characteristics of “anti” saccades in man. Exp Brain Res 89: 415–424. 1992.

Franklin DW, Wolpert DM. Specificity of reflex adaptation for task-relevant variability. J Neurosci 28: 14165–14175. 2008.

Goodale MA, Pelisson D, Prablanc C. Large adjustments in visually guided reaching do not depend on vision of the hand or perception of target displacement. Nature 320: 784–750. 1986.

Gottlieb J, Goldberg ME. Activity of neurons in the lateral intraparietal area of the monkey during an antisaccade task. Nat Neurosci 2: 906–912. 1999.

Gu C, Wood DK, Gribble PL, Corneil BD. A trial-by-trial window into sensorimotor transformations in the human motor periphery. J Neurosci 36: 8273–8282.

Häger-Ross C, Cole KJ, Johansson RS. Grip-force responses to unanticipated object loading: load direction reveals body- and gravity-referenced intrinsic task variables. Exp Brain Res 110: 142–150. 1996.

Halekoh U, Højsgaard S. A Kenward-Roger Approximation and Parametric Bootstrap Methods for Tests in Linear Mixed Models – The R Package pbkrtest. J Stat Softw 59: 1–30. 2014.

Hsu YC, Yeh CI, Huang JJ, Hung CH, Hung CP, Pei YC. Illusory motion reversal in touch. Front Neurosci 13: 605. 2019.

Hunter JS. Matplotlib: a 2D graphics environment. Comput Sci Eng 9: 90–95. 2007.

Johansson RS, Westling G. Programmed and triggered reactions to rapid load changes during precision grip. Exp Brain Res 71: 72–86. 1988.

Kenward MG, Roger JH. Small sample inference for fixed effects from restricted maximum likelihood. Biometrics 53: 983–997. 1997.

Kuznetsova A, Brockhoff PB, Christensen RHB. lmerTest Package: Tests in Linear Mixed Effects Models. J Stat Softw 82: 1–26.

Lenth R. emmeans: estimated marginal means, aka Least-squares means. R package version 1.4.4. 2020.

Luke SG. Evaluating significance in linear mixed-effects models in R. Behav Res Methods 49: 1494–1502. 2017.

McKinney W. Data structures for statistical computing in python. Proc 9th Python in Science Conf: 51–56. 2010

Ohki Y, Johansson RS. Sensorimotor interactions between pairs of fingers in bimanual and unimanual manipulative tasks. Exp Brain Res 127: 43–43. 1999.

Oliphant TE. A guide to Numpy. USA: Trelgol Publishing. 2006.

Pack CC, Bensmaia SJ. Seeing and feeling motion: Canonical computations in vision and touch. PLoS Biol 13: e1002271. 2015.

Paulignan Y, MacKenzie C, Marteniuk R, Jeannerod M. Selective perturbation of visual input during prehension movements 1. The effects of changing object position. Exp Brain Res 83: 502–512. 1991.

Pei YC, Hsiao SS, Bensmaia SJ. The tactile integration of local motion cues is analogous to its visual counterpart. Proc Natl Acad Sci USA 105: 8130–8135. 2008.

Pelisson D, Prablanc C, Goodale MA, Jeannerod M. Visual control of reaching movements without vision of the limb. Exp Brain Res 62: 303–311. 1986.

Pisella L, Grea H, Tilikete C, Vighetto A, Desmurget M, Rode G, Boisson D, Rossetti Y. An ‘automatic pilot’ for the hand in the human posterior parietal cortex: toward reinterpreting optic ataxia. Nat Nerosci 3: 729–736. 2000.

Prablanc C, Martin O. Automatic control during hand reaching at undetected two-dimensional target displacements. J Neurophysiol 67: 455–469. 1992.

Pruszynski JA, King GL, Boisse L, Scott SH, Flanagan JR, Munoz DP. Stimulus-locked responses on human arm muscles reveal a rapid neural pathway linking visual input to arm motor output. Eur J Neurosci 1049–1057. 2010.

Pruszynski JA, Johansson RS. Edge-orientation processing in first-order tactile neurons. Nat Neurosci 17: 1404–1409. 2014.

Pruszynski JA, Johansson RS, Flanagan JR. A rapid tactile-motor reflex guides reaching toward handheld objects. Curr Biol 26: 788–792. 2016.

Pruszynski JA, Flanagan JR, Johansson RS. Fast and accurate edge orientation processing during object manipulation. eLife 7:e31200. 2018.

Saijo N, Murakami I, Nishida S, Gomi H. Large-field visual motion directly induces an involuntary rapid manual following response. J Neurosci 25: 4941–4951. 2005.

Veerman MM, Brenner E, Smeets JBJ. The latency for correcting a movement depends on the visual attribute that defines the target. Exp Brain Res 187: 219–228. 2008.

Virtanen et al. SciPy 1.0: fundamental algorithms for scientific computing in Python. Nat Methods 17: 261–272. 2020.

